# MetaboRx: R Shiny application for quality assessment and preprocessing of mass spectrometry-based metabolomics data

**DOI:** 10.64898/2026.11.16.767162

**Authors:** Breanna K. Guppy, Ronald A. Merrill, Reid F. Brown, T. Idil Apak Evans, Eric B. Taylor

**Affiliations:** Department of Molecular Physiology and Biophysics, University of Iowa Carver College of Medicine, Iowa City, IA 52240, USA; Fraternal Order of Eagles Diabetes Research Center (FOEDRC), University of Iowa Carver College of Medicine, Iowa City, IA 52240, USA; FOEDRC Metabolomics Core Research Facility, University of Iowa Carver College of Medicine, Iowa City, IA 52240, USA; Holden Comprehensive Cancer Center, University of Iowa Carver College of Medicine, Iowa City, IA 52240, USA; Abboud Cardiovascular Research Center, University of Iowa Carver College of Medicine, Iowa City, IA 52240, USA; Pappajohn Biomedical Institute, University of Iowa Carver College of Medicine, Iowa City, IA 52240, USA

**Keywords:** Metabolomics, Mass Spectrometry, Software, Workflow, Normalization, Quality Control, Quality Assurance

## Abstract

**Introduction:** Metabolomics datasets offer deep insights into biological systems, yet the multi-step analytical workflow, from sample preparation through instrumental analysis, introduces unwanted variation. This variation, including signal drift and batch effects can confound true biological signals. Although rigorous quality control (QC) and data preprocessing are prerequisites for downstream analysis, existing workflows are often fragmented, time-consuming, and inaccessible to non-experts.

**Objectives:** To develop a user-friendly software tool for rapid quality assessment and rigorous preprocessing of metabolomics feature tables.

**Methods:** MetaboRx is an R Shiny application that integrates orthogonal quality metrics into a single workflow, enabling systematic and transparent evaluation of metabolomics data before and after preprocessing. The application identifies formatting errors and analytically unreliable features and provides flexible preprocessing and visualization options.

**Results:** MetaboRx is freely available at https://github.com/ETaylorLab/MetaboRx. Using a semi-targeted LC-MS metabolomics dataset from mouse plasma, perturbed with simulated low-quality data, we demonstrate how MetaboRx facilitates detection of data quality problems, reduces unwanted variation, and prepares data for downstream analysis.

**Conclusion:** MetaboRx streamlines metabolomics data quality assessment and preprocessing through an interactive and integrated workflow. The application may improve consistency and reproducibility in metabolomics studies by supporting informed preprocessing decisions and transparent reporting of QC outcomes prior to downstream analysis.

## 1 Introduction

Metabolomics, the quantification of small molecules in biological samples, is a powerful approach for studying biological systems (Clish, 2015; Dalamaga, 2024; Qiu *et al*., 2023). Liquid chromatography-mass spectrometry (LC-MS) is the most widely used platform, due to its versatility in targeted and untargeted analyses and sensitivity to a wide range of metabolites (Alanazi, 2025; Cajka, 2024; Plumb *et al*., 2023; Schrimpe-Rutledge *et al*., 2016). Nonetheless, the multi-step LC-MS workflow, from sample preparation through instrumental analysis, introduces unwanted variation that can confound true biological signals. Differences in sample preparation and evolving instrument conditions during long analytical runs propagate to shifts in retention time and peak intensity, commonly referred to as signal drift (Lu *et al*., 2017; Märtens *et al*., 2023). After metabolite identification and quantification, these technical artifacts can remain in the dataset and degrade downstream analysis.

Due to these challenges, careful data quality control and preprocessing are essential prerequisites for reliable analysis and interpretation. Without appropriate correction of technical variation, downstream analyses may be biased or difficult to interpret (Yue *et al*., 2022). Multiple computational methods have been developed to aid in the removal of non-biological variation and are implemented in different software tools. However, the effectiveness of a given method often depends on dataset-specific characteristics, including study design and the nature of technical variation. Commonly used tools such as MetaboAnalyst (Pang *et al*., 2024), NOREVA (Fu *et al*., 2022), statTarget (Luan *et al*., 2018), QuantyFey (Aigensberger *et al*., 2025), MRMkit (Teo *et al*., 2020), QC:MXP (Broadhurst *et al*., 2018), and MetaboDrift (Thonusin *et al*., 2017) provide robust correction and normalization methods, along with data visualization features. More complex and advanced methods are available with software tools such as TIGER (Han *et al*., 2022), hRUV (Kim *et al*., 2021), SERRF (Fan *et al*., 2019), and mzQuality (van der Peet *et al*., 2025). The currently available tools are excellent resources for correcting signal drift and normalizing data but provide fewer metrics for assessing data quality or selecting appropriate preprocessing methods. This leaves the users responsible for vetting quality before and after preprocessing and determining which methods are appropriate for their data.

More readily accessible, transparent data quality checks throughout the preprocessing workflow can improve the efficiency of data evaluation and the reliability of preprocessed data. Here we present MetaboRx, a user-friendly R Shiny application designed to integrate data quality assessment and preprocessing within a unified workflow. The application employs multiple quality metrics to support transparent evaluation and cleaning of metabolomics datasets. Using a mouse plasma profiling dataset perturbed by introduction of low-quality data, we show how MetaboRx identifies common data quality issues and supports preprocessing decisions. The browser-based interface dynamically reacts to the user input, accommodating both exploratory analysis and structured filtering. By combining transparent quality assessment with flexible preprocessing options, MetaboRx streamlines the preparation of metabolomics data and improves the reliability of the resulting datasets.

## 2 Materials and Methods

### 2.1 Evaluating data quality with MetaboRx

MetaboRx is implemented in R using the R Shiny framework making the app accessible to users without programming experience. The workflow guides users through quality assessment, preprocessing, evaluating, and exporting analysis-ready metabolomics data (Fig. 1). Importantly, although we focus on LC-MS metabolomic data in this manuscript, MetaboRx works equally well for data produced by GC-MS and other platforms. All outputs created by the workflow including data files, figures, and a report summarizing the process, are exported in the last module. The open-source code, installation instruction, and quick-start tutorial are all available at https://github.com/ETaylorLab/MetaboRx.

**Fig. 1.**
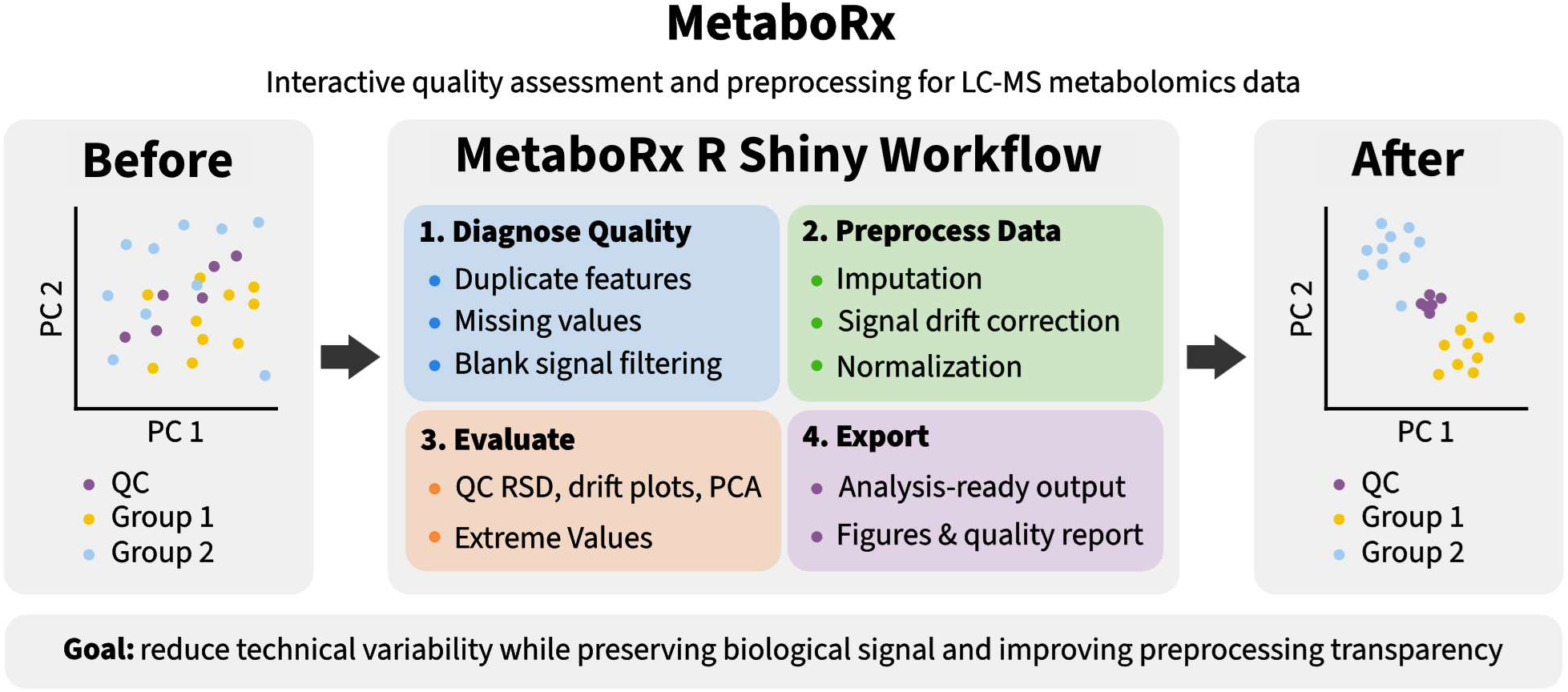
Overall workflow of LC-MS metabolomics quality assessment and preprocessing with MetaboRx.

The MetaboRx workflow aims to remove technical variation while preserving biological signal. It first corrects signal drift, then applies a sample-based normalization method as applicable. Pooled quality control (QC) samples are a widely used, technically sound approach for monitoring analytical stability and guiding signal drift correction (Broeckling *et al*., 2023; Cajka, 2024; Evans *et al*., 2020; González-Domínguez *et al*., 2024). Therefore, MetaboRx requires QC samples to be distributed throughout the run, and it uses QC-based methods for signal drift correction and variation filtering.

Signal drift correction in MetaboRx is implemented using one of four methods: local constant, local linear, local polynomial regression, or random forest regression. Local constant regression was implemented using the Nadaraya–Watson kernel regression estimator. The local linear and polynomial approaches are extensions of the commonly used QC-based robust locally weighted scatterplot smoothing signal correction (QC-RLSC) method. Like QC-RLSC, they model intensity on a per-metabolite basis as a function of injection order using the QC samples, then apply the fitted model to the experimental samples (Dunn *et al*., 2011). The random forest regression method is adapted from QC-based random forest signal correction (QC-RFSC), where random forest models are trained on QC samples using injection order as a predictor (Luan *et al*., 2018). In MetaboRx, three random forest models are trained using different random seeds, and the final corrected values are obtained by taking the median prediction across models to improve robustness. Following correction, each metabolite is multiplied by a constant so that the median corrected intensity among valid, non-imputed QC observations equals either 1 or the median of its finite, positive QC intensities before imputation and correction. The latter option retains the original QC and experimental sample intensity scale making it useful for absolute quantification methods for estimating metabolite concentration. Users may also select a “no correction” option, which omits drift modeling while allowing metabolites to be scaled to a QC median of 1 or retained on their original intensity scale.

Beyond QC normalization and drift correction, MetaboRx provides sample-based normalization options, including total ratiometric normalization (TRN) and internal standard normalization. TRN, also known as total sum normalization, first gives each QC-corrected metabolite value equal weight by scaling its mean intensity across all samples to equal one. It then scales each sample to have identical total intensities for the sum of all metabolites detected in all samples (Sun and Xia, 2024), similar in concept to total useful signal normalization (Warrack *et al*., 2009). When a control is designated, metabolite values are further scaled so that each metabolite mean in the control group equals one while preserving relative error for all metabolites in all groups. As a complement to signal-based normalization, internal standard normalization scales metabolite measurements by the average of the internal standards within each sample (De Livera *et al*., 2015). Both approaches should be used with caution since extreme values can artificially deflate other values within the same sample (Sun and Xia, 2024) and internal standards manifest their own unique variation that cannot be generalized to all measured metabolites (Sysi-Aho *et al*., 2007). To combat these issues, we include probabilistic quotient normalization as an alternative (Dieterle *et al*., 2006). Probabilistic quotient normalization corrects for differences in sample dilution. For each sample, every metabolite is divided by its median across all study samples. The median of these ratios is taken as the sample’s dilution factor, and the sample is divided by it.

MetaboRx was implemented with the assistance of Open AI Codex. OpenAI Codex was used to assist in debugging, refactoring, and optimizing code. The final code used in MetaboRx was reviewed, tested, and modified by the authors when necessary. Claude by Anthropic was used to refine and edit some of the accompanying text. We remain responsible for the software design, implementation decisions, final code, and manuscript text.

### 2.2 Mouse fasting and refeeding experiment to generate plasma for metabolomic analysis

All animal work was performed in accordance with the University of Iowa Animal Use and Care Committee (IACUC) guidelines, protocol 3101249. To test and illustrate MetaboRx’s capabilities, we designed a mouse fasting and refeeding experiment to generate plasma samples with both large and subtle metabolomic differences between groups. Sixty-four mice were assigned to eight groups of eight (age, sex, strain, and feeding regimen in Table 1). Mice were either fed normal chow diet (NCD; Teklad 2920X) ad libitum or fasted for 16h overnight. They were then either kept on ad libitum feeding or refed for 3h with NCD or a high-fat, high-sugar Western Diet (WD; Research Diets D12079Bi). Plasma was collected from every mouse at the end of the 16h fast and again after 3h of refeeding (or at the matching time points for ad libitum-fed mice), yielding 128 samples for LC-MS analysis.

**Table 1.**
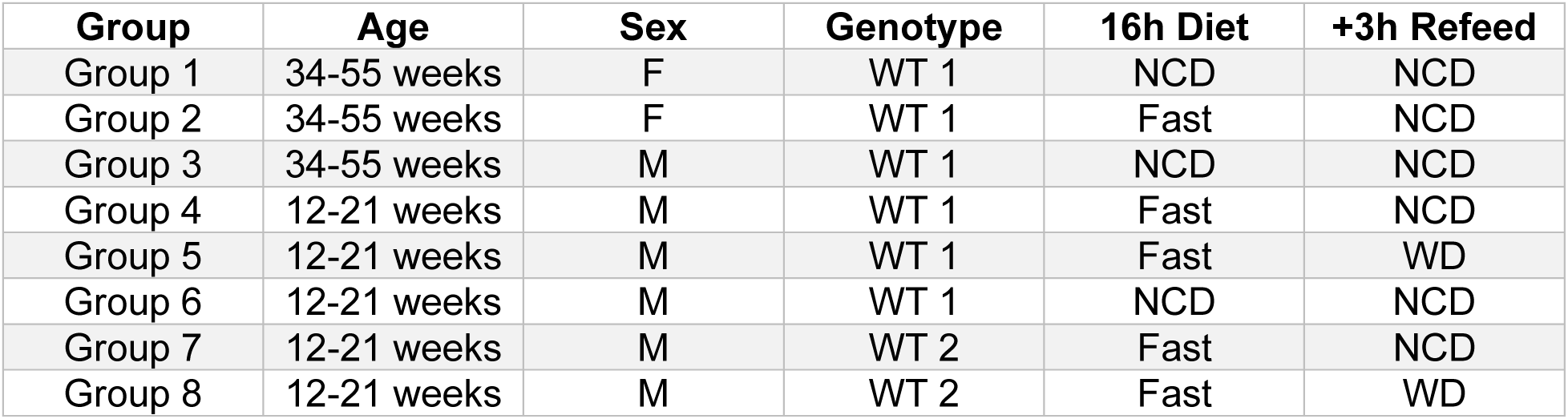
Mouse groups contain eight mice each. Genotype wild-type 1 (WT 1) and wild-type 2 (WT 2) are two different in-house mouse lines. For the 16h diet, mice were either fed normal chow diet (NCD; Teklad 2920X) ad libitum or fasted for 16h overnight (Fast). For the +3h refeed, mice either kept on ad libitum feeding or refed for 3hr with NCD or a high-fat, high-sugar Western Diet (WD; Research Diets D12079Bi).

### 2.3 Sample preparation and LC-MS analysis

For metabolite extraction, plasma samples were combined with extraction solvent (2:2:1 acetonitrile:methanol:water, v/v/v final concentration) and rotated for 1 hour at -20 °C. Processing blanks consisted of extraction solvent only and were processed identically to all other samples. Following extraction, samples were centrifuged for 10 minutes, briefly vortexed (10-15 seconds), and centrifuged again for 10 minutes. The supernatant was then collected and transferred to a new 1.5 mL tube. Pooled quality control (QC) samples were prepared by combining equal volumes of each sample extract and processed alongside study samples. Extracts were dried using a Speedvac Vacuum concentrator then resuspended in 1:1 acetonitrile:water, vortexed and incubated at -20°C overnight. Prior to analysis, samples were centrifuged, briefly vortexed, and centrifuged again before transferring the supernatant to autosampler vials. A stock human plasma sample was processed alongside study samples and used to test system suitably at the beginning of the LC-MS analysis run sequence.

LC-MS data were acquired using a Thermo Q Exactive Classic hybrid quadrupole Orbitrap mass spectrometer coupled to a Vanquish Flex UHPLC system. Chromatographic separation was performed using a SeQuant ZIC-pHILIC column (2.1 × 150 mm, 5 µm) with an 8 µL injection volume and a flow rate of 0.150 mL/min. Mobile phase A consisted of 20 mM ammonium carbonate and 0.1% ammonium hydroxide, and mobile phase B consisted of acetonitrile. The gradient decreased from 80% to 20% B over 20 minutes before returning to 80% B for column re-equilibration (Cantor et al., 2017). Pooled QC samples were injected at the beginning and end of the analytical run and after every eight study samples. Mass spectrometry was performed in full-scan polarity-switching mode over an m/z range of 70-1,000 at a resolution of 70,000, with an AGC target of 1 × 10⁶ and maximum injection time of 200 ms (Cantor *et al*., 2017). Detailed sample preparation protocol, LC-MS analysis, and run sequence are provided in the supplement (S1 Supplementary file 1).

Acquired LC-MS data were processed by Thermo Scientific TraceFinder 5.2 software, and metabolites were identified based on matching to the University of Iowa Metabolomics Core facility spectral library generated by running physical standards in house.

## 3 Results and Discussion

### 3.1 Transparent quality metrics

The experimental dataset generated as described in table 1 was exported from TraceFinder with samples arranged in rows and metabolite peak intensities in columns. Before importing the dataset into MetaboRx, sample metadata columns were added. Required metadata consists of sample class, which identifies QCs, blanks, and study sample groups, and injection order of each sample. Additional sample metadata can be included but are not required. For this dataset we include diet, sex, genotype, and age to further explore differences associated with these variables. To demonstrate MetaboRx’s signal drift correction, we introduced simulated drift into the example dataset by multiplying a random subset of metabolites by injection order-dependent scaling factors. The supplementary file includes the original and perturbed data, the affected metabolites and their scaling factors, and the MetaboRx-formatted raw data (S2 Supplementary file 2).

Missing values are common in metabolomics data, which makes filtering features by missingness a critical step in any analysis workflow. MetaboRx first removes metabolites that are all zero or missing across all QC and study samples. In the example dataset, HMG-CoA, malonyl-CoA, and propionyl-CoA, which are typically not detected in plasma, were intentionally included in the search list to demonstrate this feature (Fig. 2a). Metabolites are then filtered using separate user-defined missing value thresholds for study samples and QCs, each with a default value of 20%, consistent with the QComics workflow (González-Domínguez *et al*., 2024). Users can choose between two criteria for removing metabolites, depending on the sample classes being analyzed. For the first, a metabolite can be removed if its percentage of missing values exceeds the designated threshold in any sample class. For the second, metabolites are removed only if the threshold is exceeded in all sample classes. The second option is appropriate when group missingness may reflect real biological differences. For ease of evaluation and transparency, removed metabolites are listed under the failed threshold along with any remaining metabolites with missing QC values (Fig. 2b). Notably, here, the six retained metabolites with missing QC values are expected to be very low abundance in plasma and potentially originated from cell lysis during plasma preparation. Users can bypass missing value filtering by increasing the thresholds to 100% while still receiving warnings when metabolites are entirely absent for a specific sample class. Missing value frequencies in metabolites, samples, batches, and classes offer additional analytical and biological insights and are available for download.

**Fig. 2.**
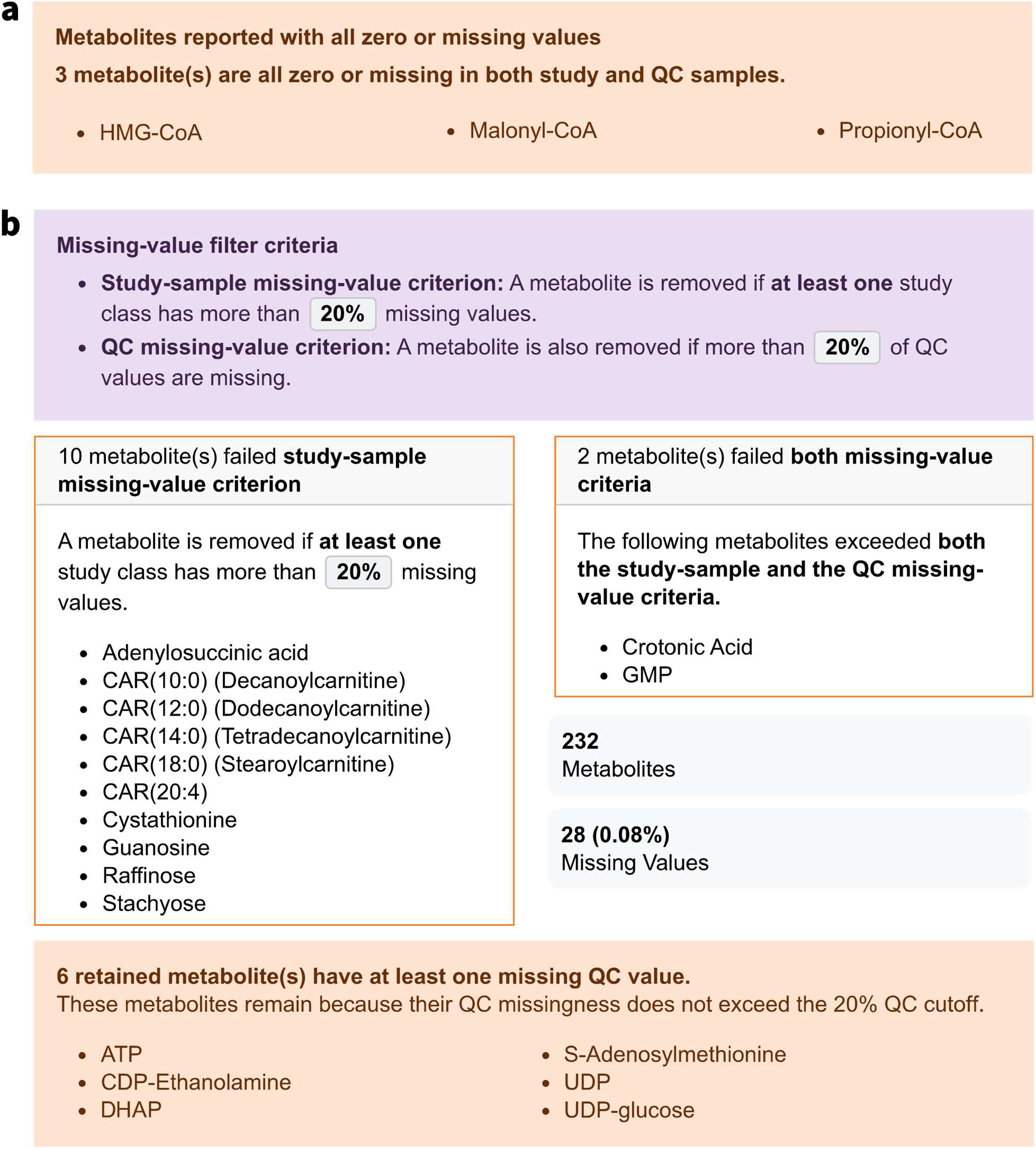
MetaboRx missing value filtering. (a) Metabolites with all zero or missing values are removed from the dataset during missing value filtering. We intentionally included these metabolites here to test this feature. (b) Missing value summary at the default missing value thresholds including missing value counts and percentages after filtering.

After correction, metabolites are evaluated based on their relative standard deviation (RSD; standard deviation divided by mean), of QC samples, an accepted measure for analytical reproducibility in metabolomics (Parsons *et al*., 2009). Metabolites exceeding the 30% QC RSD threshold suggested by mQACC are removed and listed for inspection (Kirwan *et al*., 2022). Thresholds are user-adjustable and both pre- and post-correction RSD values are available for download. Users can inspect the changes in RSD using the plots generated by MetaboRx (Fig. 3a). Additionally, warnings notify the user if spike-in labeled internal standards exceed the QC RSD threshold and if a metabolite’s study sample average differs from QC average by a user set percentage of the QC mean (Fig. 3b-c). These features summarize the quality of the QC-based correction ensuring reliable metabolite measurements proceed to downstream analysis.

**Fig. 3.**
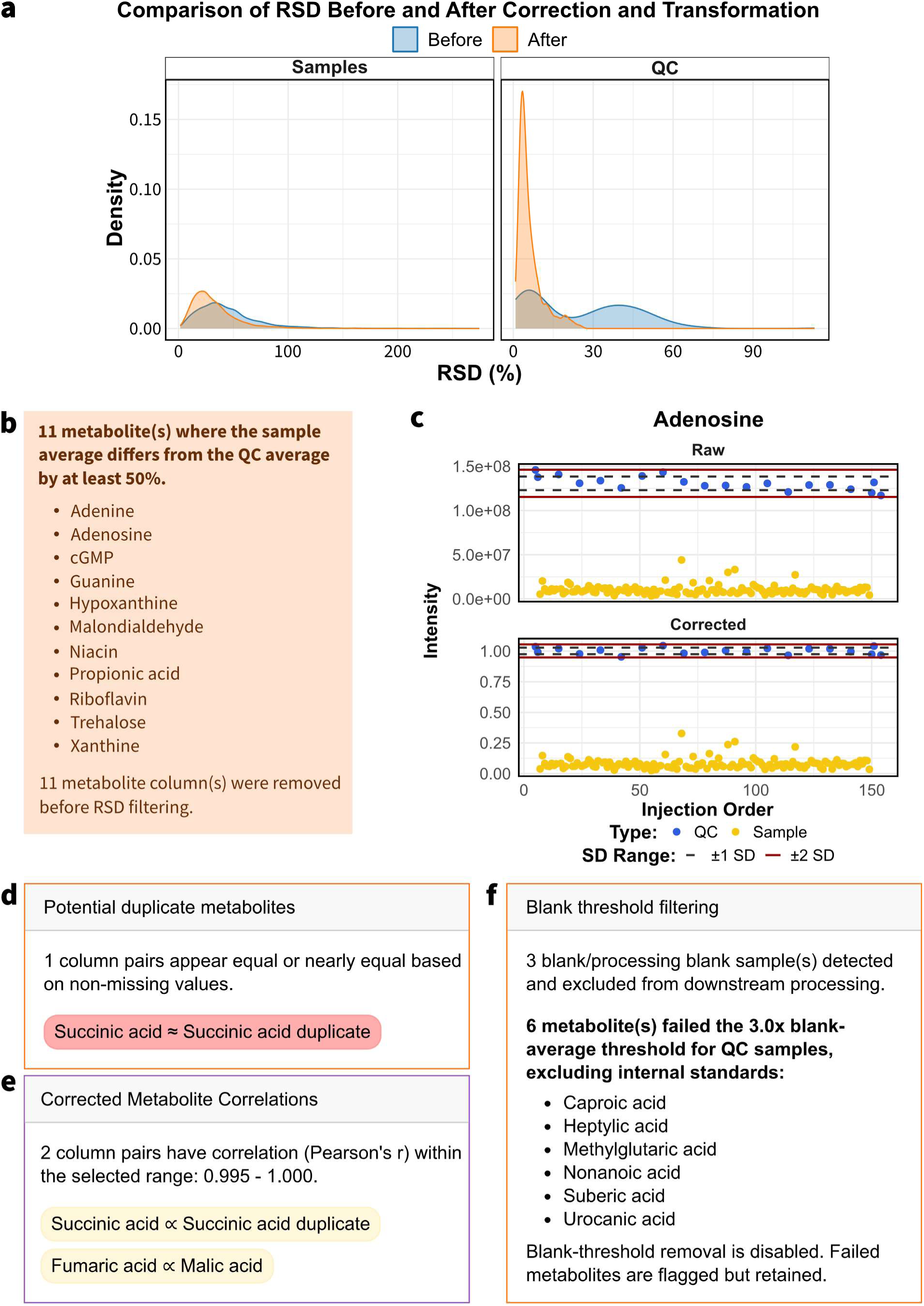
Quality filtering metrics. (a) Sample RSD computed within class and QC RSD distributions before and after correction and TRN. Density on the y-axis indicates distribution of RSD values with a smooth curve. (b) Warning from MetaboRx for metabolites with a sample average that differs from QC average by at least 50% of the QC average, potentially reflecting in sample chemistry related to additional QC handling steps. (c) Scatter plot of adenosine intensity before and after correction illustrating the large difference in sample vs QC average. (d) Potential duplicate metabolite warning based on equal values. (e) Highly correlated metabolites indicating possible duplicate metabolites as well as expected biological correlations in the signal drift corrected data. (f) Blank threshold filtering summary flagging potential contaminates or very low abundance metabolites.

Distinguishing isobaric and isomeric metabolites remains a challenge necessitating the removal of incorrect or redundant measurements (Ben Faleh *et al*., 2023; Opialla *et al*., 2020). MetaboRx addresses this issue by flagging metabolites with equal or strong positively correlated values. Strong positively correlated metabolites are displayed in the application window, and all pairwise correlations are available for download. To demonstrate this feature, we duplicated succinic acid to show that MetaboRx will warn users of the duplicate by checking for equal values (Fig. 3d) and flagging highly correlated metabolites (Fig. 3e). Some metabolites may exhibit biologically driven correlations because they participate in closely linked metabolic reactions. For example, fumaric acid and malic acid are directly interconverted through the reversible fumarase-catalyzed reaction, providing a biological basis for correlated abundances between these metabolites. Due to the complexity of correctly identifying which metabolites are truly detected in the samples, duplicates are only flagged, and the choice of removal is left to the user.

Processing blanks or true blanks, when available, are used to identify potential contamination or background signal. MetaboRx uses blanks to flag metabolites whose QC average intensity is less than 3 times the average blank intensity, as suggested by the metabolomics quality assurance and quality control consortium (mQACC) (Evans *et al*., 2020). This threshold is user-adjustable, and flagged metabolites can be optionally excluded. For the example dataset, metabolites that fail the blank threshold are possible contaminates or have very low abundance in study samples (Fig. 3f).

Extreme values exert a disproportionate influence on total metabolite load and some sample-based normalization techniques (Warrack *et al*., 2009). Their exclusion reduces variability at the risk of removing important biological information (Playdon *et al*., 2019). MetaboRx identifies candidate extreme values using two complementary approaches. Potential outlier samples are detected using Mahalanobis distances in principal component analysis (PCA) two-dimensional score space, with a chi-squared threshold analogous to Hotelling’s *T*^2^ described in the QComics workflow (González-Domínguez *et al*., 2024) (Fig. 4a). In lay terms, any study sample that sufficiently stands out from all other samples in the simplified PCA space is flagged as an outlier. Independently, log transformed metabolite intensities are screened for extreme values within each sample class. For each metabolite, sample classes with fewer than eight observations were tested using a two-sided Grubbs-type test (α = 0.0005) based on externally studentized residuals from one-way ANOVA. The residual variance was pooled across classes for that metabolite after omitting the observation being tested, and p-values were Bonferroni-adjusted within each class (Fox and WeisBerg, 2019; Grubbs, 1969). Classes with eight or more observations were tested using an intercept-only adaptation of ROUT (Q = 0.01%) (Motulsky and Brown, 2006) (Fig. 4b). These extreme value thresholds are included to bring awareness to potential outliers, but the decision for formal exclusion is left to investigators and their statisticians according to best practices. This approach prioritizes identification of extreme values while minimizing unnecessary exclusion, particularly in small datasets.

**Fig. 4.**
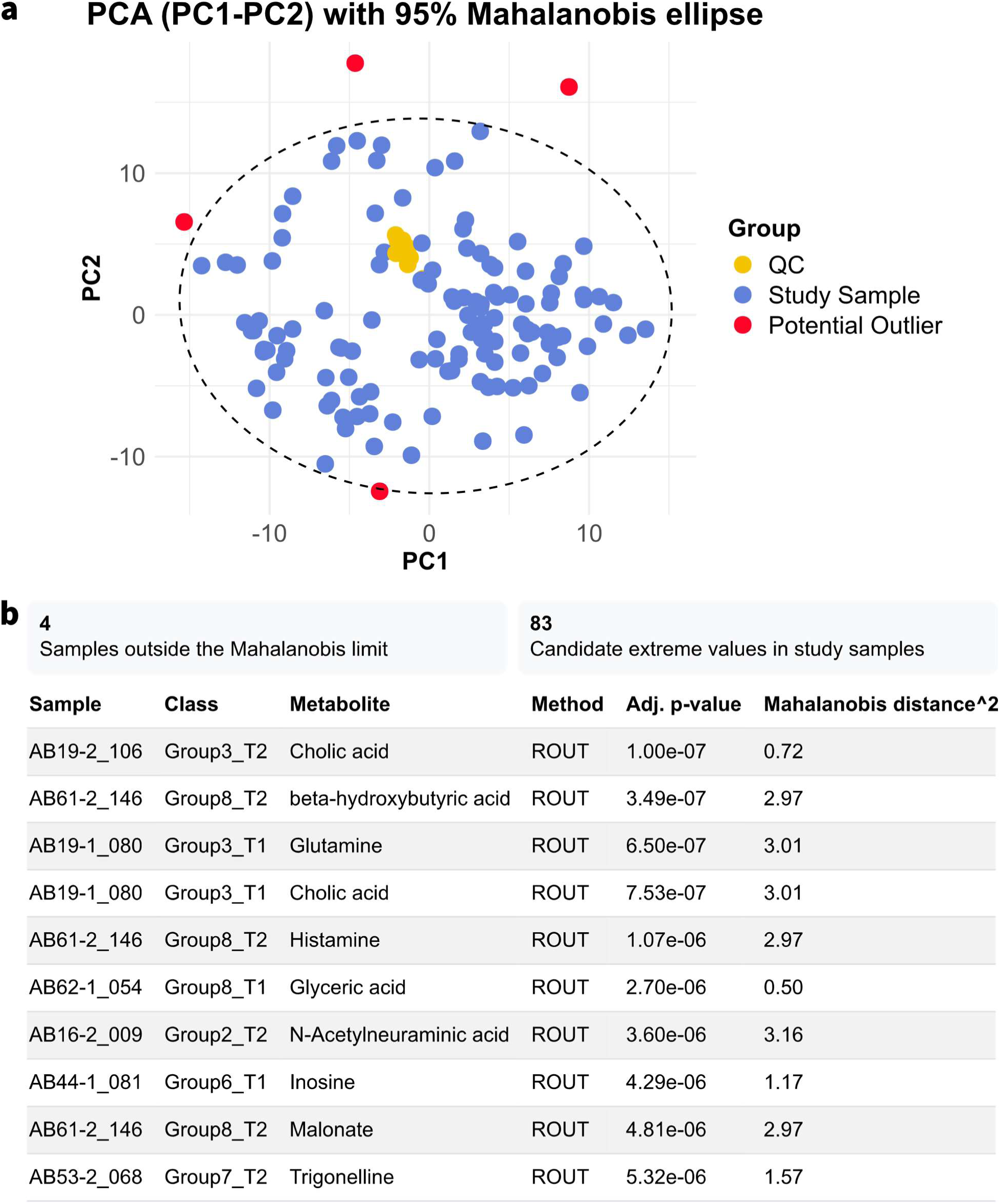
Potential extreme value detection. (a) PCA score plot with Mahalanobis 95% ellipse with outlier samples colored red. (b) Potential extreme value summary and table displaying top 10 extreme values sorted by adjusted p-value within MetaboRx.

### 3.2 Adaptable options

Since QC-based regression models rely on QC sample intensities and are sensitive to outliers, especially when the number of QC samples is limited, missing values must be imputed prior to correction (Motulsky and Brown, 2006). MetaboRx allows users to apply different imputation strategies for QC and study samples, enabling greater flexibility in handling missing values. Available imputation methods include minimum value (Li *et al*., 2017), mean, kNN (Troyanskaya *et al*., 2001), median, half the minimum value, and zero imputation (Armitage *et al*., 2015). These imputation methods use the information provided in the non-missing data to estimate what a missing value should be. Because imputation can bias downstream analyses, particularly metabolite correlation structure (Gromski *et al*., 2014; Taylor *et al*., 2017), imputed values are marked by highlighting in the tabulated output. Alternatively, users can select “remove imputed values” after correction, resulting in blank cells instead. If imputed values are removed, the imputed values will still have been used for PCA plots.

Signal drift patterns vary across datasets in both scale and complexity, making the selection of an appropriate correction method nontrivial. Previous studies have shown that no single method consistently outperforms others across all datasets (De Graeve *et al*., 2023; Liao *et al*., 2023; Märtens *et al*., 2023). In general, what correction method is best for a run depends on its size and the complexity of the drift patterns (Thonusin *et al*., 2017). To support correction method selection, MetaboRx provides guidance based on the total number of QCs and their spacing within the run (Fig. 5a). More complex methods, such as random forest, require a greater number of well-distributed QC samples compared to simpler regression-based approaches. Therefore, the random forest method is only available when there is at least a total of 9 QC samples in the dataset with no more than 10 study samples in between QCs.

**Fig. 5.**
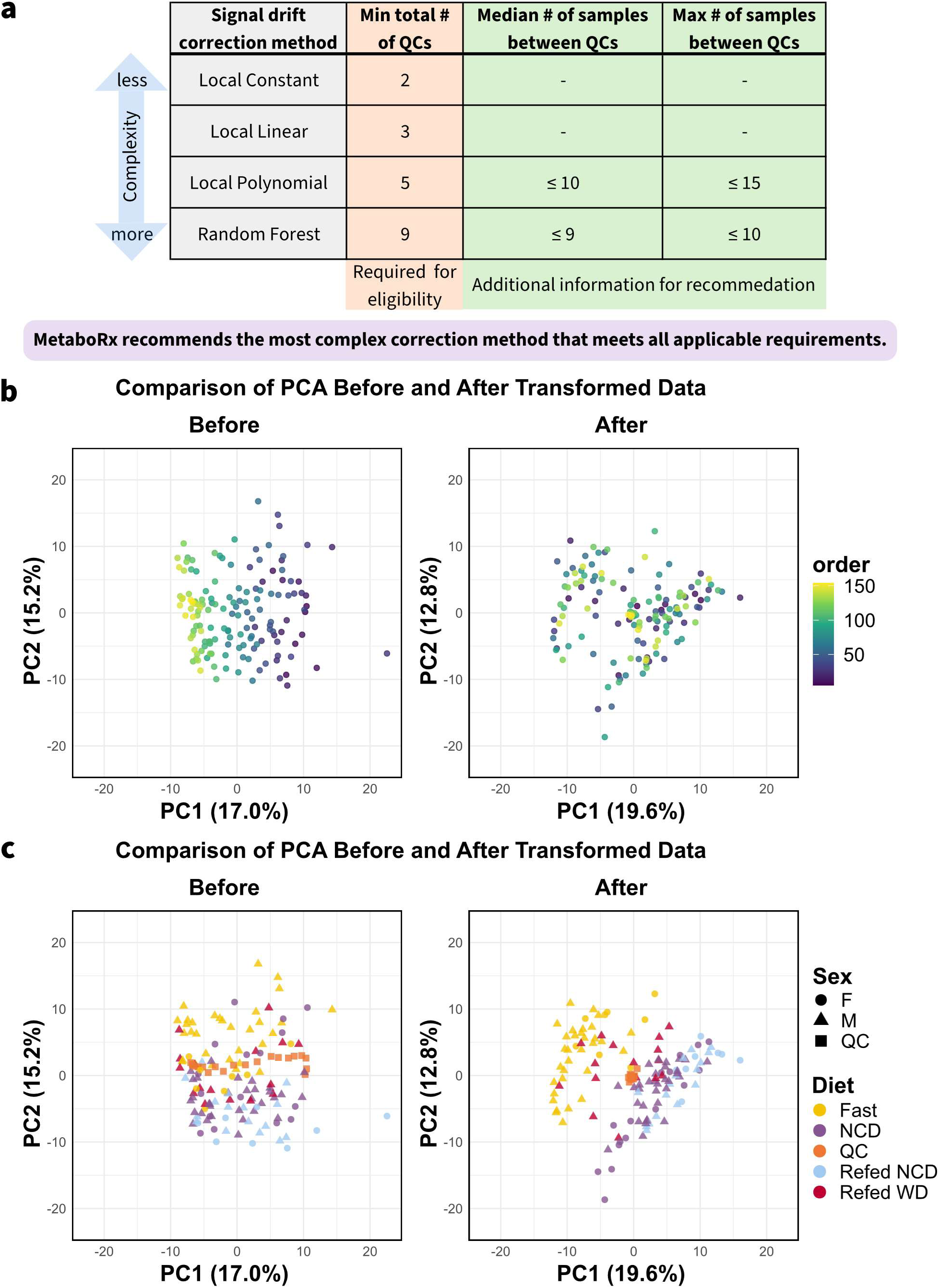
MetaboRx has adaptable options and flexible visualization setting. (a) MetaboRx’s framework for correction method eligibility and recommendation. Suggested correction method is based on the total number of QCs in the experiment as well as the frequency of QCs in the run. (b) PCA score plot of the perturbed experimental dataset with simulated signal drift before and after correction and sample normalization colored by injection order. (c) PCA score plot of the perturbed experimental dataset with simulated signal drift before and after correction and sample normalization colored by diet with shapes determined by sex.

MetaboRx offers multiple visualization tools to assess data quality and preprocessing performance. Scatter plots of metabolite intensity versus injection order compare raw and corrected data, facilitating visual evaluation of the signal drift correction method. RSD plots summarize changes in analytical variability before and after correction. PCA score plots provide an overview of sample structure and can be colored by user-specified metadata variables, such as diet or injection order, to aid interpretation of biological and technical variation. In the perturbed dataset, coloring the PCA plot by injection order highlights the simulated signal drift in the raw data and its removal in corrected data (Fig. 5b). In addition to coloring, secondary metadata can be used to select the shape of the sample’s point. In the perturbed dataset utilized here, samples are better clustered by diet post-correction and normalization (Fig. 5c). In addition to visualization, MetaboRx generates downloadable figures, summary reports, and formatted datasets, supporting efficient transition to downstream statistical analysis.

## 4 Conclusions

Metabolomics studies generate large, complex datasets that require preprocessing and quality assessment before biological interpretation. While numerous software tools exist for signal drift correction, normalization, and statistical analysis, fewer resources focus on guiding users through quality assessment and preprocessing decisions. MetaboRx addresses this need by integrating quality assessment and preprocessing into a single workflow, emphasizing transparency and informed decision-making throughout the data cleaning process.

MetaboRx provides an integrated framework for metabolomics data quality assessment and preprocessing, combining multiple quality metrics with flexible, user-driven processing options. By embedding transparent evaluation steps throughout the workflow, the application enables users to systematically diagnose data quality issues, evaluate preprocessing effectiveness, and make informed decisions prior to downstream analysis. Through features such as missing value assessment, QC-based filtering, signal drift correction diagnostics, extreme value detection, duplicate value detection, and general metabolite correlation analysis, MetaboRx consolidates tasks that are often distributed across multiple software platforms.

MetaboRx provides numerous advantages, but its limitations should also be acknowledged. Although MetaboRx was developed and demonstrated using LC-MS data, the workflow can also be applied to other mass spectrometry-based metabolomics datasets, including gas chromatography (GC-MS) and ion mobility spectrometry (IMS-MS), provided that the data are formatted appropriately and include pooled QC samples. The effectiveness of the workflow also depends on experimental design and data quality, as no preprocessing strategy can fully compensate for poor sample preparation, insufficient QC coverage, or severe analytical variability. Furthermore, while the application provides guidance and quality metrics to assist decision-making, the final choice of preprocessing methods and filtering thresholds remains dependent on user judgment and study-specific objectives.

The primary goal of MetaboRx is to make rigorous metabolomics data preprocessing more accessible while enabling consistent and reproducible quality assessment practices. By combining complementary quality checks, interactive visualizations, flexible preprocessing options, and transparent reporting within a single application, MetaboRx lowers the technical barriers associated with metabolomics data cleaning. We anticipate that this framework will facilitate more informed preprocessing decisions, improve transparency in data handling, and support the generation of higher-quality datasets for downstream biological interpretation.

## Supplementary Information

S1 Supplement file 1: Full detailed sample preparation and LC-MS protocol.

S2 Supplement file 2: Excel workbook containing the original and perturbed example datasets, the metabolites affected by simulated drift and their scaling factors, the MetaboRx-formatted raw data, and the sample information and LC-MS run sequence.

## Acknowledgements

We thank Paulina Sobieralski for assisting during the plasma collection experiment and Satya Murthy Tadinada for feedback on software usability.

## Author Contributions

BKG, RAM, and EBT conceived the project, developed the software, performed data analysis, designed the experiment. BKG and EBT wrote and edited the manuscript. BKG designed, implemented, validated, documented, and reviewed the code. BKG and RAM performed the experiment. RFB, TIAE, RAM, EBT, and BKG reviewed and evaluated the application during development.

## Data Availability

The raw metabolomics data reported in this paper are available at the NIH Common Fund’s National Metabolomics Data Repository (NMDR) website, the Metabolomics Workbench, https://www.metabolomicsworkbench.org (Sud *et al*., 2016) under study identifier ST005427.

## Software Availability

The software developed in this study is available via https://github.com/ETaylorLab/MetaboRx.

## Declarations

### Ethical Statements

All animal work was performed in accordance with the University of Iowa Animal Use and Care Committee (IACUC) guidelines, protocol 3101249.

