## Supplementary material for "MetaboRx: R Shiny application for quality assessment and preprocessing of mass spectrometry-based metabolomics data": S1 Supplementary file 1

**Supplement 1**

1. **Sample preparation and LC-MS analysis**

For metabolite extraction, 10µL plasma samples were diluted in 680µL of ice-cold 2:2:1 methanol/acetonitrile/water where the 1-part water was composed of sample volume + water. The extraction buffer additionally contained a mixture of internal standards (D4-citric acid, D4-succinic acid, D8-valine, and U^13^C-labeled glutamine, glutamic acid, lysine, methionine, serine, and tryptophan; Cambridge Isotope Laboratories), at a concentration of 1 µg/ml each. The samples were then inverted to mix and incubated while gently rotating at -20°C for 1 hour. After incubating, the samples were centrifuged for 10 minutes at maximum speed (21100g), briefly vortexed (10-15 seconds), and centrifuged again for 10 minutes. The supernatants were then transferred to fresh 1.5 ml microcentrifuge. A pooled quality control (QC) sample was prepared by adding an equal volume of each sample to a fresh 1.5 ml microcentrifuge tube. Processing blanks were utilized by adding extraction solvent to microcentrifuge tubes. Then a 80µL aliquot of all samples, pooled QCs, and processing blanks were evaporated in 1.5 ml microcentrifuge tubes using a speed-vac at room temperature. A human plasma sample was prepared alongside study samples and used to test system suitably at the beginning of the LC-MS analysis run sequence.

Dried extracts were reconstituted in 40 µL acetonitrile/water (1:1 v/v), vortexed well, and incubated in a -20C freezer overnight. In the morning, the resuspended samples were vortexed and centrifuged for 10 minutes at maximum speed, and the supernatant was transferred to LC-MS autosampler vials for analysis.

LC-MS data were acquired on a Thermo Q Exactive hybrid quadrupole Orbitrap mass spectrometer with a Vanquish Flex UHPLC system. The LC column used was a Millipore SeQuant ZIC-pHILIC (2.1 × 150 mm, 5 µm particle size) with a ZIC-pHILIC guard column (20 × 2.1 mm). The injection volume was 8 µL. Two mobile phases were used, solvent “A” containing 20 mM ammonium carbonate [(NH4)2CO3] and 0.1% Ammonium Hydroxide (v/v) [NH4OH] [Note: pH is ~9.1] and solvent “B” containing 100% Acetonitrile. The method was run at a flow rate of 0.150 mL/min.

The gradient started at 80% B and decreased to 20% B over 20 minutes before returning to 80% B in 0.5 minutes and held there for 7 minutes (Cantor *et al.*, 2017). Note: there is a 2-minute pre-equilibration time prior to sample injection and during re-equilibration from 24.5 to 26.5 minutes flow is increased to 0.3 ml/min. The pooled QC samples were injected at the beginning, end, and every 8 samples throughout the run.

The mass spectrometer was operated in full-scan, polarity-switching mode from 1 to 20 minutes, with the spray voltage set to 3.0 kV, the heated capillary held at 275 °C, and the HESI probe held at 350 °C. The sheath gas flow was set to 40 units, the auxiliary gas flow was set to 15 units, and the sweep gas flow was set to 1 unit. MS data acquisition was performed in a range of m/z 70-1,000, with the resolution set at 70,000, the AGC target at 1 × 10^6^, and the maximum injection time at 200 ms (Cantor *et al.*, 2017).

2. **Run Sequence**

| **Sample name** | **Injection Order** | **Sample Class** |
| --- | --- | --- |
| Blank1 | 1 | blank |
| PB | 2 | processing blank |
| HP | 3 | human plasma |
| Blank2 | 4 | blank |
| QC01 | 5 | QC |
| QC02 | 6 | QC |
| AB28-2 | 7 | study sample |
| AB04-1 | 8 | study sample |
| AB16-2 | 9 | study sample |
| AB56-1 | 10 | study sample |
| AB43-1 | 11 | study sample |
| AB64-2 | 12 | study sample |
| AB39-2 | 13 | study sample |
| AB22-1 | 14 | study sample |
| QC03 | 15 | QC |
| AB35-1 | 16 | study sample |
| AB15-2 | 17 | study sample |
| AB26-2 | 18 | study sample |
| AB44-2 | 19 | study sample |
| AB53-1 | 20 | study sample |
| AB63-1 | 21 | study sample |
| AB01-1 | 22 | study sample |
| AB24-2 | 23 | study sample |
| QC04 | 24 | QC |
| AB14-2 | 25 | study sample |
| AB56-2 | 26 | study sample |
| AB18-2 | 27 | study sample |
| AB41-1 | 28 | study sample |
| AB51-1 | 29 | study sample |
| AB34-1 | 30 | study sample |
| AB31-2 | 31 | study sample |
| AB06-1 | 32 | study sample |
| QC05 | 33 | QC |
| AB57-1 | 34 | study sample |
| AB02-1 | 35 | study sample |
| AB48-2 | 36 | study sample |
| AB17-1 | 37 | study sample |
| AB13-1 | 38 | study sample |
| AB36-2 | 39 | study sample |
| AB62-2 | 40 | study sample |
| AB29-2 | 41 | study sample |
| QC06 | 42 | QC |
| AB27-2 | 43 | study sample |
| AB21-1 | 44 | study sample |
| AB55-1 | 45 | study sample |
| AB42-2 | 46 | study sample |
| AB52-2 | 47 | study sample |
| AB01-2 | 48 | study sample |
| AB39-1 | 49 | study sample |
| AB10-1 | 50 | study sample |
| QC07 | 51 | QC |
| AB23-2 | 52 | study sample |
| AB07-2 | 53 | study sample |
| AB62-1 | 54 | study sample |
| AB45-2 | 55 | study sample |
| AB38-2 | 56 | study sample |
| AB12-1 | 57 | study sample |
| AB58-1 | 58 | study sample |
| AB29-1 | 59 | study sample |
| QC08 | 60 | QC |
| AB25-2 | 61 | study sample |
| AB45-1 | 62 | study sample |
| AB38-1 | 63 | study sample |
| AB10-2 | 64 | study sample |
| AB18-1 | 65 | study sample |
| AB63-2 | 66 | study sample |
| AB05-2 | 67 | study sample |
| AB53-2 | 68 | study sample |
| QC09 | 69 | QC |
| AB27-1 | 70 | study sample |
| AB12-2 | 71 | study sample |
| AB60-2 | 72 | study sample |
| AB61-1 | 73 | study sample |
| AB46-2 | 74 | study sample |
| AB40-1 | 75 | study sample |
| AB06-2 | 76 | study sample |
| AB24-1 | 77 | study sample |
| QC10 | 78 | QC |
| AB50-2 | 79 | study sample |
| AB19-1 | 80 | study sample |
| AB44-1 | 81 | study sample |
| AB33-1 | 82 | study sample |
| AB32-2 | 83 | study sample |
| AB54-1 | 84 | study sample |
| AB11-2 | 85 | study sample |
| AB07-1 | 86 | study sample |
| QC11 | 87 | QC |
| AB05-1 | 88 | study sample |
| AB14-1 | 89 | study sample |
| AB59-2 | 90 | study sample |
| AB25-1 | 91 | study sample |
| AB41-2 | 92 | study sample |
| AB50-1 | 93 | study sample |
| AB33-2 | 94 | study sample |
| AB21-2 | 95 | study sample |
| QC12 | 96 | QC |
| AB30-1 | 97 | study sample |
| AB54-2 | 98 | study sample |
| AB23-1 | 99 | study sample |
| AB49-1 | 100 | study sample |
| AB15-1 | 101 | study sample |
| AB37-2 | 102 | study sample |
| AB04-2 | 103 | study sample |
| AB43-2 | 104 | study sample |
| QC13 | 105 | QC |
| AB19-2 | 106 | study sample |
| AB09-2 | 107 | study sample |
| AB60-1 | 108 | study sample |
| AB03-2 | 109 | study sample |
| AB49-2 | 110 | study sample |
| AB36-1 | 111 | study sample |
| AB42-1 | 112 | study sample |
| AB32-1 | 113 | study sample |
| QC14 | 114 | QC |
| AB51-2 | 115 | study sample |
| AB09-1 | 116 | study sample |
| AB58-2 | 117 | study sample |
| AB28-1 | 118 | study sample |
| AB22-2 | 119 | study sample |
| AB46-1 | 120 | study sample |
| AB35-2 | 121 | study sample |
| AB03-1 | 122 | study sample |
| QC15 | 123 | QC |
| AB64-1 | 124 | study sample |
| AB30-2 | 125 | study sample |
| AB37-1 | 126 | study sample |
| AB08-2 | 127 | study sample |
| AB48-1 | 128 | study sample |
| AB20-2 | 129 | study sample |
| AB11-1 | 130 | study sample |
| AB55-2 | 131 | study sample |
| QC16 | 132 | QC |
| AB13-2 | 133 | study sample |
| AB34-2 | 134 | study sample |
| AB52-1 | 135 | study sample |
| AB17-2 | 136 | study sample |
| AB59-1 | 137 | study sample |
| AB26-1 | 138 | study sample |
| AB47-2 | 139 | study sample |
| AB08-1 | 140 | study sample |
| QC17 | 141 | QC |
| AB16-1 | 142 | study sample |
| AB47-1 | 143 | study sample |
| AB31-1 | 144 | study sample |
| AB02-2 | 145 | study sample |
| AB61-2 | 146 | study sample |
| AB40-2 | 147 | study sample |
| AB20-1 | 148 | study sample |
| AB57-2 | 149 | study sample |
| QC18 | 150 | QC |
| QC19 | 151 | QC |
| ddNeg_QC19 | 152 | ddms Negative QC |
| ddPos_QC20 | 153 | ddms Positive QC |
| QC20 | 154 | QC |
